# Glucocorticoid receptor collaborates with pioneer factors and AP-1 to execute genome-wide regulation

**DOI:** 10.1101/2021.06.01.444518

**Authors:** Erin M. Wissink, Delsy M. Martinez, Kirk T. Ehmsen, Keith R. Yamamoto, John T. Lis

## Abstract

The glucocorticoid receptor (GR) regulates transcription through binding to specific DNA motifs, particularly at enhancers. While the motif to which it binds is constant across cell types, GR has cell type-specific binding at genomic loci, resulting in regulation of different genes. The presence of other bound transcription factors (TFs) is hypothesized to strongly influence where GR binds. Here, we addressed the roles of other TFs in the glucocorticoid response by comparing changes in GR binding and nascent transcription at promoters and distal candidate cis-regulatory elements (CCREs) in two distinct human cancer cell types. We found that after glucocorticoid treatment, GR binds to thousands of genomic loci that are primarily outside of promoter regions and are potentially enhancers. The majority of these GR binding sites are cell-type specific, and they are associated with pioneer factor binding. A small fraction of GR occupied regions (GORs) displayed increased bidirectional nascent transcription, which is a characteristic of many active enhancers, after glucocorticoid treatment. Non-promoter GORs with increased transcription were specifically enriched for AP-1 binding prior to glucocorticoid treatment. These results support a model of transcriptional regulation in which multiple classes of TFs are required. The pioneer factors increase chromatin accessibility, facilitating the binding of GR and additional factors. AP-1 binding poises a fraction of accessible sites to be rapidly transcribed upon glucocorticoid-induced GR binding. The coordinated activity of multiple TFs then results in cell type-specific changes in gene expression. We anticipate that many models of inducible gene expression also require multiple distinct TFs that act at multiple steps of transcriptional regulation.

## Introduction

Transcription of gene networks can be regulated by a variety of extracellular stimuli, including developmental signals, hormones, and environmental changes. This regulation is specified by DNA sequence elements associated with promoters and enhancers (Zabidi, Stark 2016; Long et al. 2016). These regulatory elements bind to sequence-specific transcription factors (TFs) that can alter DNA accessibility, bring in co-activators, and recruit the pre-initiation complex, including RNA Polymerase II (Pol II) (Core, Adelman 2019). While gene promoters are required for transcription, they are often not sufficient and instead additionally require the activity of distal enhancers (Zabidi, Stark 2016).

Enhancers are sequences that regulate target promoters by acting as platforms for TF binding (Zabidi, Stark 2016; Long et al. 2016). These elements, also termed response elements (Chandler et al. 1983; Weikum et al. 2017), are located within or between genes, do not necessarily regulate the nearest promoter, and can associate with candidate target promoters via direct looping (Schoenfelder, Fraser 2019), apparently enabling enhancers to be located a megabase or more from their target (Lettice et al. 2002). While genes are frequently transcribed in many cell types, enhancers generally have more cell-type specific activity because they bind TFs or other co-factors that have restricted expression and/or assemble factors in combinatorial fashion based on cell-specific conditions (Andersson, Gebhard, et al. 2014; Weikum et al. 2017).

Enhancer activities are commonly ascribed to, or correlated with, chromatin accessibility and histone modifications, particularly H3K27ac and more H3K4me1 than H3K4me3 (Heintzman et al. 2007; Ernst, Kellis 2010). However, assessing which enhancers have regulatory activity, and defining the molecular features that determine that activity, remains an area of open investigation. One model predicts that accessible sites with H3K27ac that are in contact with promoters are active (Fulco et al. 2019). Others have shown that those distal sites that are transcribed as enhancer RNAs (eRNAs) (Ashe et al. 1997; De Santa et al. 2010; Kim et al. 2010; D Wang, Garcia-Bassets, et al. 2011) have greater functional activity in reporter assays than non-transcribed sites (Henriques et al. 2018; Mikhaylichenko et al. 2018; Wissink et al. 2019). These eRNAs can be detected with high sensitivity by sequencing nascent RNAs because they produce bidirectional transcripts, similar to promoters (Andersson, Gebhard, et al. 2014; Core, Martins, et al. 2014; Scruggs et al. 2015; Mikhaylichenko et al. 2018; Wissink et al. 2019). The architecture of promoters and enhancers are similar, as TFs bind in the nucleosome-depleted region between the two core promoters for both types of elements (Core, Martins, et al. 2014; Andersson, Sandelin, Danko 2015; Tome et al. 2018; Tippens et al. 2020), and the specific TFs bound likely distinguish enhancers from promoters (Andersson, Sandelin 2020).

The TFs that bind to regulatory elements play different roles, and the constellation of TFs that can bind determine the stimuli to which enhancers respond (Weikum et al. 2017; Vihervaara, Duarte, et al. 2018). An individual enhancer can bind to multiple TFs, including ones that bind to closed chromatin (“pioneer factors”), chromatin remodelers, and factors that release Pol II from promoter-proximal pausing (Andersson, Sandelin 2020). The well-studied heat shock system demonstrates the importance of multiple factors acting at different steps of transcription. HSF1 activates transcription of hundreds of genes by binding to the DNA motif known as the heat shock element, then releasing paused polymerases into active elongation (Bunch et al. 2014; Mahat, Salamanca, et al. 2016). However, less than half of the HSF1-bound promoters have increased transcription after heat shock (Mahat, Salamanca, et al. 2016), demonstrating that this TF is not sufficient for transcription. Instead, activation of genes by HSF1 relies on pre-binding of other factors (Duarte et al. 2016), accessible chromatin (Vihervaara, Mahat, et al. 2017), and pre-formed enhancer-promoter loops (Ray et al. 2019). Similarly, signaling from inflammation uses pre-wired looping (Jin et al. 2013) and pre-binding of a pioneer factor (Escoubet-Lozach et al. 2011). Thus, these studies of transcription networks have uncovered general principles of how other factors impact gene regulation, at least for stress-related genes.

In the current work, we investigated transcriptional regulation by the glucocorticoid receptor (GR). Like HSF1, GR binds to a DNA motif, in this case the GR binding sequences (GBSs), in response to an extracellular stimulus, namely, glucocorticoid hormones. These ligands bind to cytoplasmic GR, triggering its release from chaperones, and allowing it to translocate to the nucleus to bind GBSs (Miranda et al. 2013; Meijsing 2015; Weikum et al. 2017). GR recognizes a cell type-specific subset of GBS motifs in different cell types, resulting in regulation of cell type-specific target genes (So et al. 2007; John et al. 2011; Love, Huska, et al. 2017; Thormann et al. 2018; Franco et al. 2019). GR mostly binds to accessible chromatin (John et al. 2011; Grøntved et al. 2013; Love, Huska, et al. 2017) that is pre-marked with H3K27ac and EP300 binding (McDowell et al. 2018) at sites that are typically >10 kb from promoters (So et al. 2007; Reddy et al. 2009). GR binds to roughly 1 × 10^4^ specific genomic loci in a given cell type, yet only a small fraction of these GR-occupied regions (GORs) are thought to have enhancer activity (Vockley et al. 2016). While studies have interrogated the chromatin features and factors that allow GR binding, how these are coordinated to create a functional enhancer remains unclear.

Here, we are utilizing two cell types with different lineages to disentangle the roles of different TF classes in the GR response. These cell lines are A549, a lung adenocarcinoma, and U2OS, an osteosarcoma. With nascent RNA sequencing, we determined which genes and transcribed putative enhancers are commonly and specifically regulated in response to GR signaling in these cells. The transcribed putative enhancers comprise only a small fraction of the GORs in each cell type, so we identified TF motifs that are associated with transcription. Finally, we compared the cell type-specific factors that are enriched at cell type-specific GORs. With these data, we demonstrate the importance of the binding of TFs that potentially act in different roles, including making chromatin more accessible, enabling the activity of GR, and activating transcription in the cellular response to glucocorticoids, and we discuss the features common to other transcriptionally inducible systems.

## Results

### GR signaling has a stronger transcriptional effect in U2OS cells than A549 cells

We investigated GR’s transcriptional regulatory activity in two cell lines, a lung carcinoma (A549) and an osteosarcoma (U2OS). A549 cells have endogenous GR expression, whereas these U2OS cells have transgenic GR expression (Rogatsky, Trowbridge, et al. 1997). To examine the primary effects of GR on the nascent transcriptome, we treated these cells with either ethanol (control) or the synthetic corticosteroid dexamethasone (dex) for 45 minutes, followed by PRO-seq (Figure 1A). Increases in nascent transcription are readily apparent at this time point, as seen at the *FKBP5* gene, which is induced by dex in both cell types (Figure 1B). Both cell types have genes that are up- or downregulated after dex treatment, although far more genes are differentially expressed after dex treatment in U2OS than in A549 cells (Figure 1C-E, Figure S1A,B, Table S1), showing that the U2OS cells are more responsive to GR signaling. The changes in gene transcription correlate significantly between the two cell types (Figure 1E). While 450 genes that are differentially expressed in A549 cells show a concordant and significant change in expression in U2OS cells, 322 genes show A549-specific expression changes, and 2011 have U2OS-specific effects (Figure 1F).

**Figure 1:**
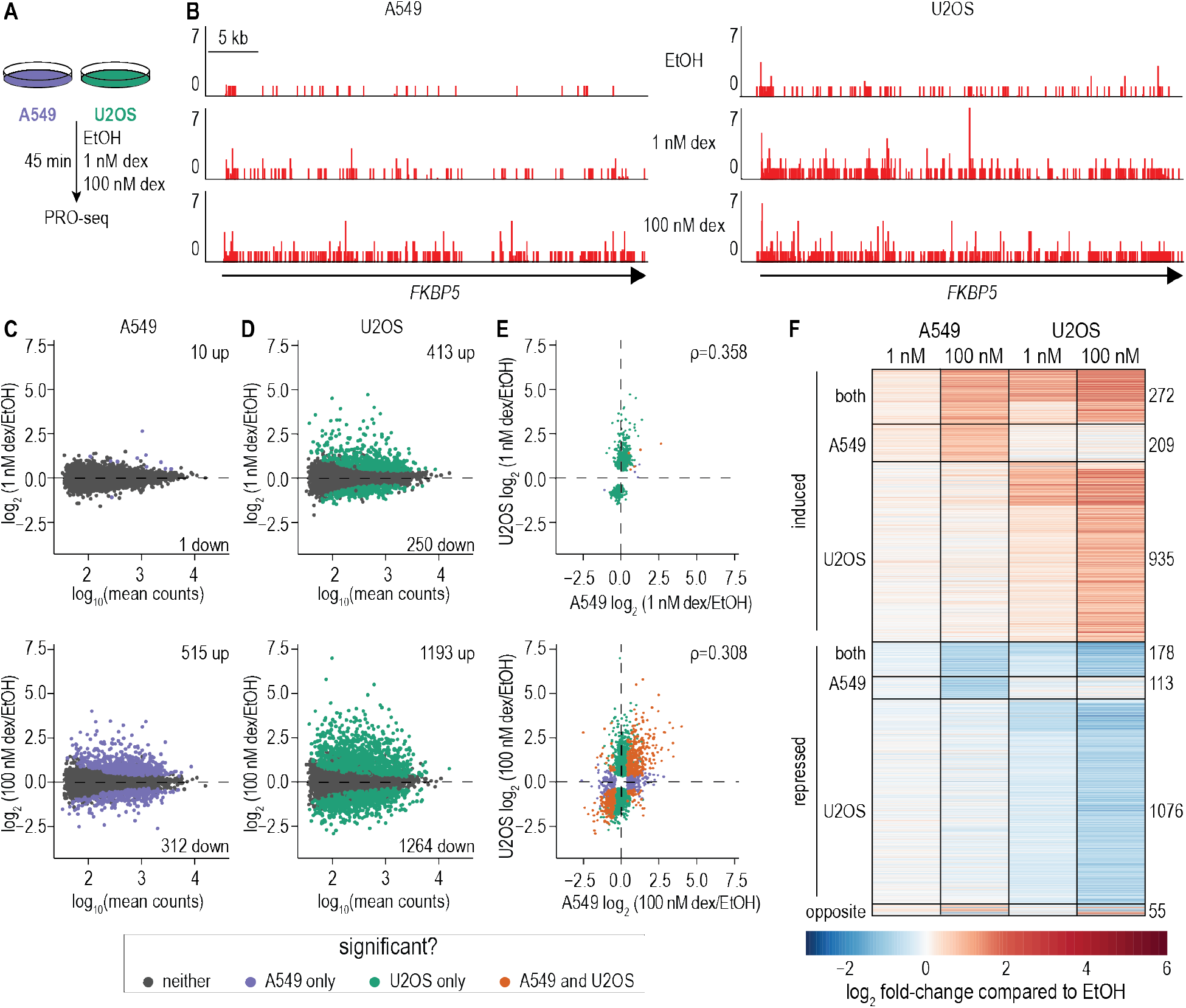
Genic transcription changes after glucocorticoid stimulation. **(A)** Experimental set-up. **(B)** Browser shot depicting nascent RNA from PRO-seq reads for the beginning of the *FKBP5* gene in A549 (left) and U2OS (right) cells. Coordinates are chr6:35,634,000-35,690,000 (hg38). **(C, D)** MA plots for global gene expression changes in A549 **(C)** and U2OS **(D)** cells, comparing treatment with ethanol (vehicle control, EtOH) to treatment with 1 nM dex (upper) or 100 nM dex (lower). Numbers of significantly differentially expressed genes are denoted; Benjamini-adjusted p < 0.05. **(E)** Comparison of fold-change differences in expression for genes called as significantly different in A549 and/or U2OS cells, after treatment with 1 nM dex (upper) or 100 nM dex (lower). Spearman correlation coefficient is displayed; p < 2.2 × 10^-16^ for both. **(F)** Expression change heat-map for genes in panel E. Genes have been categorized by their response to 100 nM dex treatment.

A previous study showed that some mature mRNAs are significantly downregulated after 3 hours of treatment with 1 nM dex but upregulated with 100 nM dex, or vice versa (Chen et al. 2013). Dose-dependent gene expression changes are not apparent in the nascent transcriptome after 45 minutes dex treatment (Figure S1C,D), suggesting this incoherent feed-forward loop occurs later in signaling and/or post-transcriptionally.

One possible explanation for the differences in GR responsiveness is that U2OS cells have 5 times more GR protein than A549 cells (Chen et al. 2013), yet the fact that A549 and U2OS cells each have unique target genes suggest that other factors are also playing a critical role. Some genes are inversely dex-responsive in the two cell types tested (i.e., activated in one and repressed in the other) (Figure 1E,F), further indicating that GR is not the sole determinant of the cellular response to dex.

### GR binds different sites in A549 and U2OS cells

To identify underlying causes of cell-type specific gene expression changes after dex treatment, we performed ChIP-seq to GR in A549 and U2OS cells after a 90 min treatment with 100 nM dex. We found thousands of GR-occupied regions (GORs) across the genome in each cell type, with more widespread occupancy in U2OS cells (Figure 2A, Table S2). Each cell type has specific binding sites, yet there are 4596 that are shared (63% of 7313 for A549, 18% of 24,891 for U2OS). We compared our ChIP-seq data to published ATAC-seq data (ENCODE Project Consortium 2012; Davis et al. 2018; BH Lee, Stallcup 2018), and found that GR binds to both highly and lowly accessible chromatin in both A549 and U2OS cells (Figure 2A).

**Figure 2:**
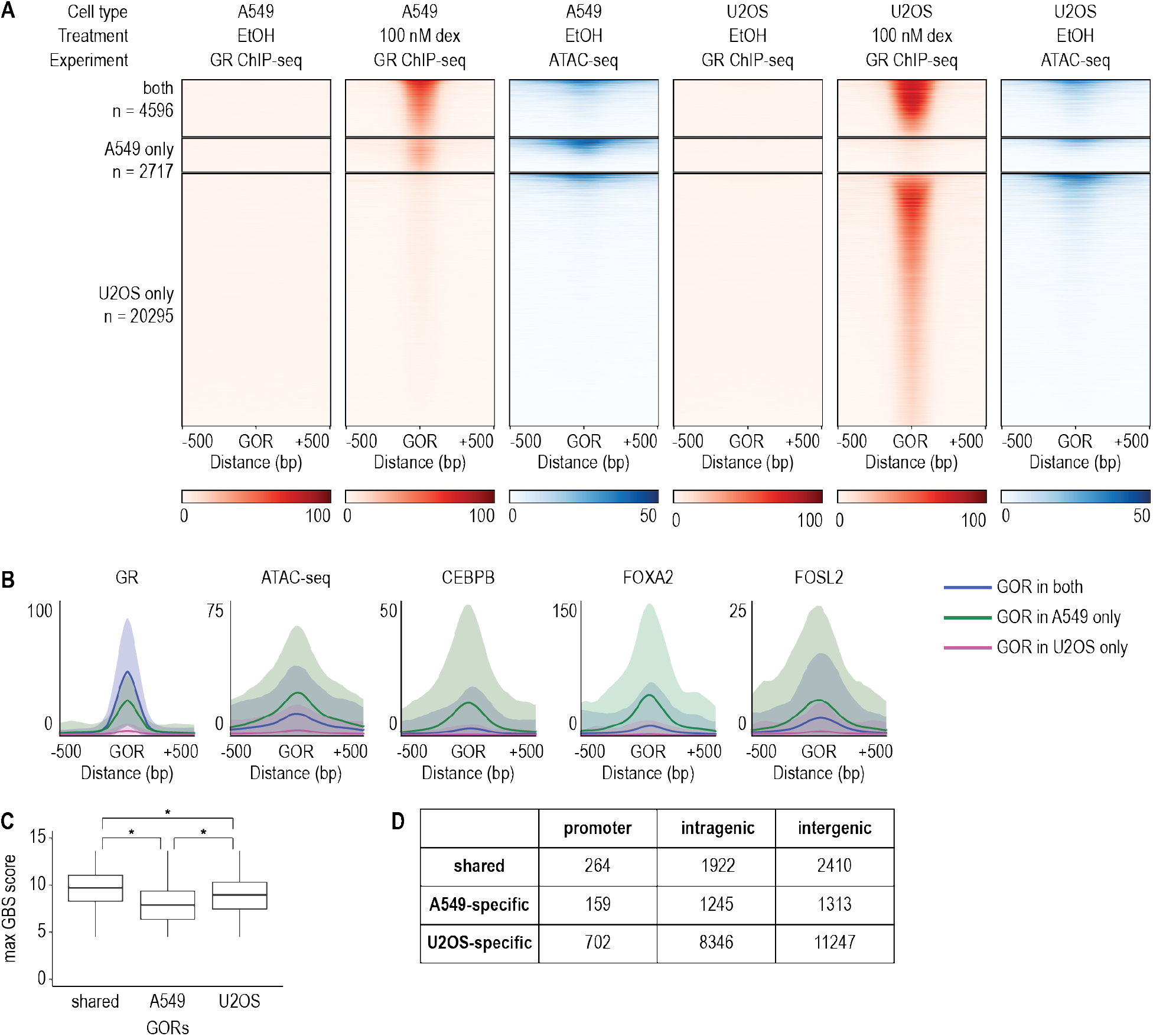
Cell type determinants of GR binding. **(A)** GR-occupied regions (GORs) were identified from GR ChIP-seq signal and categorized as being shared between A549 cells, specific to A549 cells, or specific to U2OS cells. GR ChIP-seq signal is shown in control cells and cells treated with 100 nM dex; ATAC-seq signal is shown in control cells. Signal is -log10(p-value) of enrichment. **(B)** In A549 cells, metaplots of mean signal for GR ChIP-seq in 100 nM dex-treated cells, and ATAC-seq and ChIP-seq data for CEBPB, FOXA2, and FOSL2 in control cells are shown for shared GORs (blue), A549-specific GORs (green), and U2OS-specific GORs (pink). Signal is the -log10 p-value of enrichment, and shading indicates the standard deviation. **(C)** The log-odds scores for the GR binding sequence (GBS) were found for GORs that are shared or cell-type specific. The score of the best GBS for each is plotted. * p < 1 × 10^-8^. **(D)** Genomic distribution of GORs.

The cell type-specific binding of GR demonstrates that information beyond the GBS directs its binding, so we used motif searching to identify other factors that may associate with GORs. Our search was limited to +/- 150 bp from the summit of GR binding, as this distance captures the regulatory information between two core promoters that are associated with candidate enhancers (Core, Martins, et al. 2014; Scruggs et al. 2015; Tippens et al. 2020). Motifs for FOXA, CEBP, and AP-1 (which consists of a heterodimer of FOS, JUN, and ATF family members, or c-JUN homodimers) were enriched in A549-specific GORs, as compared to U2OS-specific GORs (Table S3). Additionally, expression of FOXA1, FOXA2, CEBPA, and CEBPB are significantly upregulated in A549 cells, as compared to U2OS cells, whereas some but not all members of the AP-1 complex have higher A549 expression (Figure S2A). Each of these factors have reported pioneer activity (Lupien et al. 2008; Belikov et al. 2009; Biddie et al. 2011; Grøntved et al. 2013) and have been implicated in affecting GR binding (So et al. 2007; Belikov et al. 2009; Biddie et al. 2011; Grøntved et al. 2013; Starick et al. 2015; Vockley et al. 2016; McDowell et al. 2018; Hoffman et al. 2018). The higher relative expression of FOXA and CEBP in A549 cells may increase chromatin accessibility at A549-specific GORs, enabling GR binding.

One highly significantly enriched motif at U2OS-specific GORs, as compared to A549-specific GORs, is the RUNX motif (Table S3). The RUNX family of transcription factors has pioneering activity D Wang, Diao, et al. 2018; Veeken et al. 2019; JW Lee et al. 2019) and may act in higher-order chromatin structure (Barutcu et al. 2016). RUNX2 is required for osteoblast differentiation, is expressed in U2OS cells (Komori 2005; Lucero et al. 2013), and has significantly higher expression in U2OS than A549 cells (Figure S2A). Other motifs include the GBS (GCR) and the nearly identical progesterone receptor motif (PRGR) (Table S3). While the GBS enrichment is significant, the GBS is found at a high fraction of both U2OS- and A549-specific GORs.

Conceivably, U2OS GR binding may be enabled by the higher expression of RUNX2, which could increase chromatin accessibility at U2OS-specific GORs.

Protein-binding data are more informative than the presence of enriched motifs. Unfortunately, little publicly available ChIP-seq data for U2OS cells exist; however, ChIP-seq data have been published for FOXA2, CEBPB, FOSL2, JUN, and JUNB in A549 cells (Donaghey et al. 2018; McDowell et al. 2018). We compared the presence of these factors at GORs that are either shared or cell type-specific in cells that were not treated with dex. We also compared chromatin accessibility by ATAC-seq prior to dex treatment (ENCODE Project Consortium 2012; Davis et al. 2018; BH Lee, Stallcup 2018). Together, the binding of these factors and ATAC-seq reveal the chromatin landscape available to GR when its binding is rapidly activated by dex treatment (Figure 2B, Figure S2B). A549-specific GORs are overall the most accessible prior to dex addition, and they have the strongest binding of CEBPB, FOXA2, and AP-1 components (Figure 2B, Figure S2B), consistent with the observed motif enrichment. Sites that are specific to U2OS cells display the least accessibility and are not enriched for motifs for the A549-enriched components: CEBPB, FOXA2, and AP-1 (Figure 2B, Figure S2B). While shared GORs have less binding of these pioneer factors, they have the greatest magnitude of GR binding and higher AP-1 binding than U2OS-specific sites. These results suggest that pioneer factor-binding prior to dex treatment promotes chromatin accessibility and thereby allows GR binding to cell type-specific sites. We predict that in U2OS cells, binding of RUNX2 guides GR binding. The higher GR binding at GORs that are shared between U2OS and A549 cells suggests these sites rely on other TFs, genomic features, and/or the quality of the GBS motif.

To examine the impact of GBS strength, we found the log odds scores of every GBS at a GOR (Figure 2C). Shared GORs have the strongest GBS, followed by U2OS GORs, then A549 GORs. Similarly, estrogen receptor (ER) binding sites that are shared between two cell types have stronger ER elements than cell-type specific ones (Gertz et al. 2013). Further, GORs that are specific to A549 cells have significantly fewer GBS elements per binding site than U2OS-specific or shared sites (Figure S2C). GORs that are shared between cell types may not rely on the prior binding of a cell type-specific pioneer factor. Instead, the ubiquitously expressed AP-1 may bind and open chromatin, and then multiple GBS elements that strongly bind to GR could permit GR to destabilize nucleosome binding to chromatin.

We interrogated the genomic loci to which GR binds (Figure 22D) and characterized binding sites as being a promoter (within 1 kb from an annotated TSS), or being at intra- and intergenic sites, some of which may be enhancers. The vast majority of sites (96%) are at non-promoter loci and are the subject of the remainder of this study. We did also examine the genes with GR binding at their promoters. For the small number of promoter-associated GORs, it may prove interesting that most are linked to dex-induced, as opposed to dex-repressed genes (Figure S2D,E). However, a role for promoter-bound GR in regulation remains to be examined.

### Enhancers have cell type-specific transcriptional changes in response to GR signaling

The expression and activity of enhancers are largely cell-type specific (Andersson, Gebhard, et al. 2014), and transcribed eRNAs can be readily captured by PRO-seq (Wissink et al. 2019). Here, we used the program dREG to identify sites of bidirectional transcription (Z Wang et al. 2019; Chu et al. 2019) from our PRO-seq data, then called loci that were at least 1 kb from an annotated transcription start site as putative enhancers (candidate cis-regulatory elements, CCREs) (Moore et al. 2020). Of the 32,237 bidirectional transcripts detected in A549 and U2OS cells, 13,522 overlapped promoters, while the 11,207 intergenic and 7508 intragenic loci were considered CCREs (Figure S3A). The promoter-matching transcripts were predominantly shared between cell types, whereas CCREs were more likely to be cell type-specific (Figure S3A). We found that 5% of CCREs exhibit a significant change in transcription after dex treatment (dex-responsive CCREs; Table S4). As seen in the genic transcription, dex-responsive CCREs are largely cell type-specific, with a larger dex response in U2OS than in A549 cells (Figure 3). Examples of CCREs are shown (Figure 3A), including one that is dex-inducible in both cell types (left), in only A549 cells (center), and in only U2OS cells (right). GR binding, as measured by ChIP-seq after 100 nM dex treatment, is also included. In one of the example CCREs, GR binding is present in both cell types, yet changes in transcription are present only in A549 cells (Figure 3A, center). As discussed in more depth below, binding of GR is not sufficient to predict which CCREs will be differentially expressed in each cell type.

**Figure 3:**
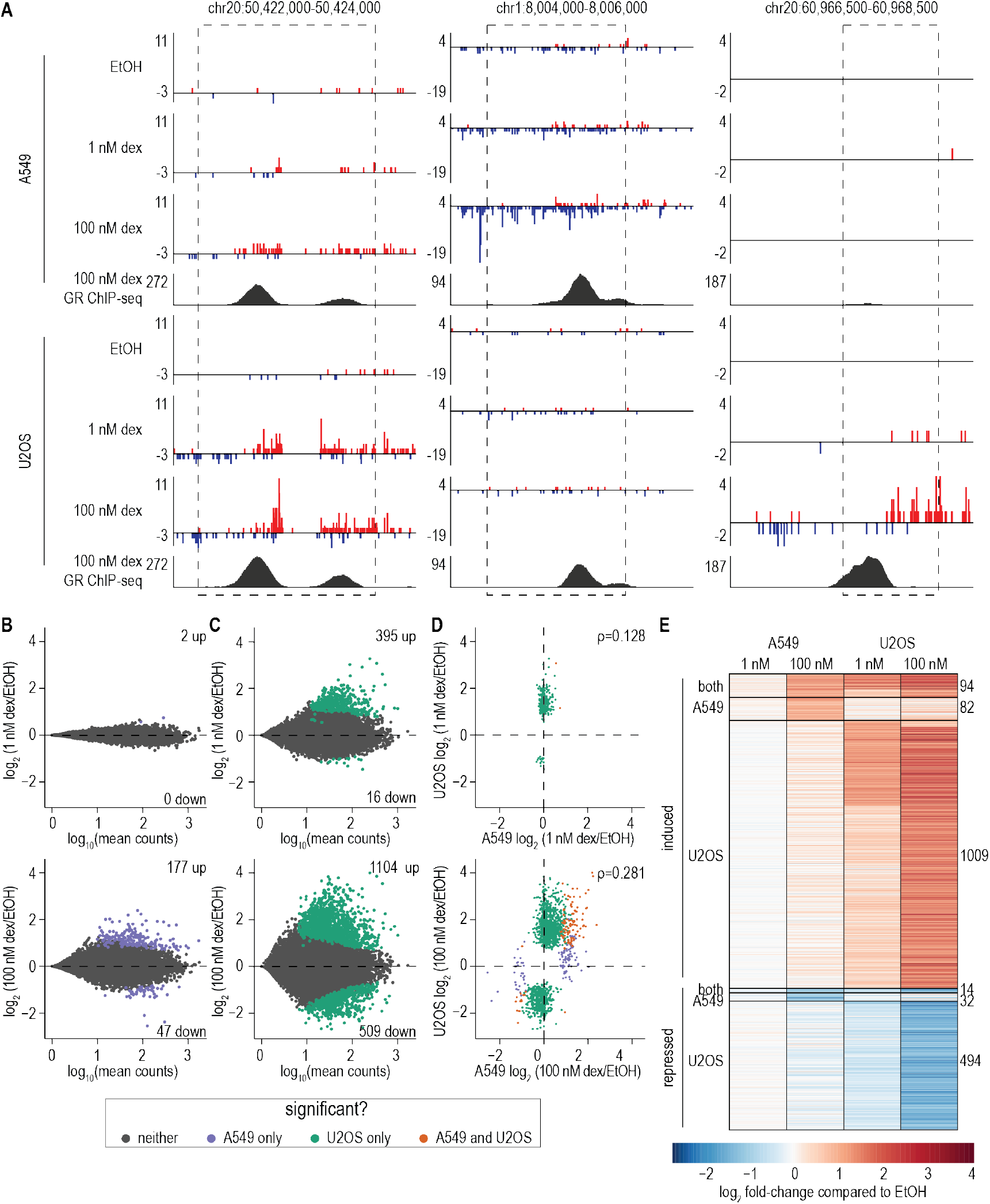
Candidate *cis*-regulatory element (CCRE) transcription changes after glucocorticoid stimulation. **(A)** Browser shot depicting nascent RNA from PRO-seq (upper three panels) and GR ChIP-seq (lower) in A549 and U2OS cells. Included are a CCRE induced in both cell types (left), only in A549 cells (center), and only in U2OS cells (right). In PRO-seq, red and blue indicate sense and anti-sense transcription, respectively. Dashed boxes delimit CCREs determined by dREG. **(B, C)** MA plots for global CCRE expression changes in A549 **(B)** and U2OS **(C)** cells, comparing treatment with EtOH to treatment with 1 nM dex (upper) or 100 nM dex (lower). Numbers of significantly differentially expressed genes are shown; Benjamini-adjusted p < 0.05. **(D)** Comparison of fold-change differences in expression for CCREs called as significantly different in A549 and/or U2OS cells, after treatment with 1 nM dex (upper) or 100 nM dex (lower). Spearman correlation coefficient is displayed; p < 2.2 × 10^-16^ for both. **(E)** Expression change heat-map for CCREs in panel **D**. CCREs have been categorized by their response to 100 nM dex treatment. One CCRE with opposing responses in A549 and U2OS cells was omitted from the heatmap.

The global trends for dex-responsive transcription at CCREs are similar to genes. A subset of CCREs have the same response in both cell types, but overall U2OS cells have greater changes in transcription at both 1 nM and 100 nM dex treatment (Figures 3B-E, S2B-D). The stronger U2OS response is apparent after the 1 nM treatment, where only two CCREs have significant differential expression in A549 cells, as compared to 411 CCREs in U2OS, and the changes between cell types have a smaller correlation than after 100 nM dex treatment (Figure 3B-E). While there are similar numbers of up- and down-regulated genes, CCREs are more likely to exhibit induced rather than repressed transcription after dex treatment in both cell types (Figures 3B-E).

### CCREs showing changes in PRO-seq signal have characteristics of GR-induced enhancers

Our identification of dex-responsive CCREs was based solely on PRO-seq data, and thus did not incorporate information from proteins binding to DNA, transcription factor motifs, histone modifications, or chromatin accessibility. We therefore looked for expected features of GR-responsive enhancers in these CCREs. Induced CCREs are enriched for the GR binding motif in both cell types (p=0.01 in A549, p=1 × 10^-26^ in U2OS, Table S3). The repressed CCREs in A549 cells are not significantly enriched in any TF motifs, while repressed CCREs in U2OS cells are enriched in motifs for homeobox factors, SMAD, CREM, and zinc finger proteins (Table S3). GR could act on repressed CCREs via tethering to the locus by another DNA binding factor (Weikum et al. 2017) or instead by an indirect mechanism.

Approximately half of enhancers appear to act on the nearest gene (Fulco et al. 2019), and so we determined if the closest expressed gene to each dex-responsive CCRE is dex-responsive. While not every closest gene was dex-responsive, nearest genes to induced CCREs have globally increased expression after dex treatment in both cell types (Figure 4A, B), genes nearest repressed CCREs are commonly repressed by dex (Figure 4C, D). As with the repressed CCREs, the repressed nearest genes displayed little binding of GR at their promoters, implying further that their repression is likely an indirect consequence of GR signaling. We also determined that dex-responsive genes are located near dex-responsive CCREs. Because a gene’s expression can be influenced by multiple enhancers, we did not necessarily expect that the nearest CCRE would be dex-responsive. In both cell types, however, genes that are upregulated are closer to induced CCREs than genes that are downregulated or unaffected by dex treatment (Figure 4E,G), while the converse is true for downregulated genes (Figure 4F,H).

**Figure 4:**
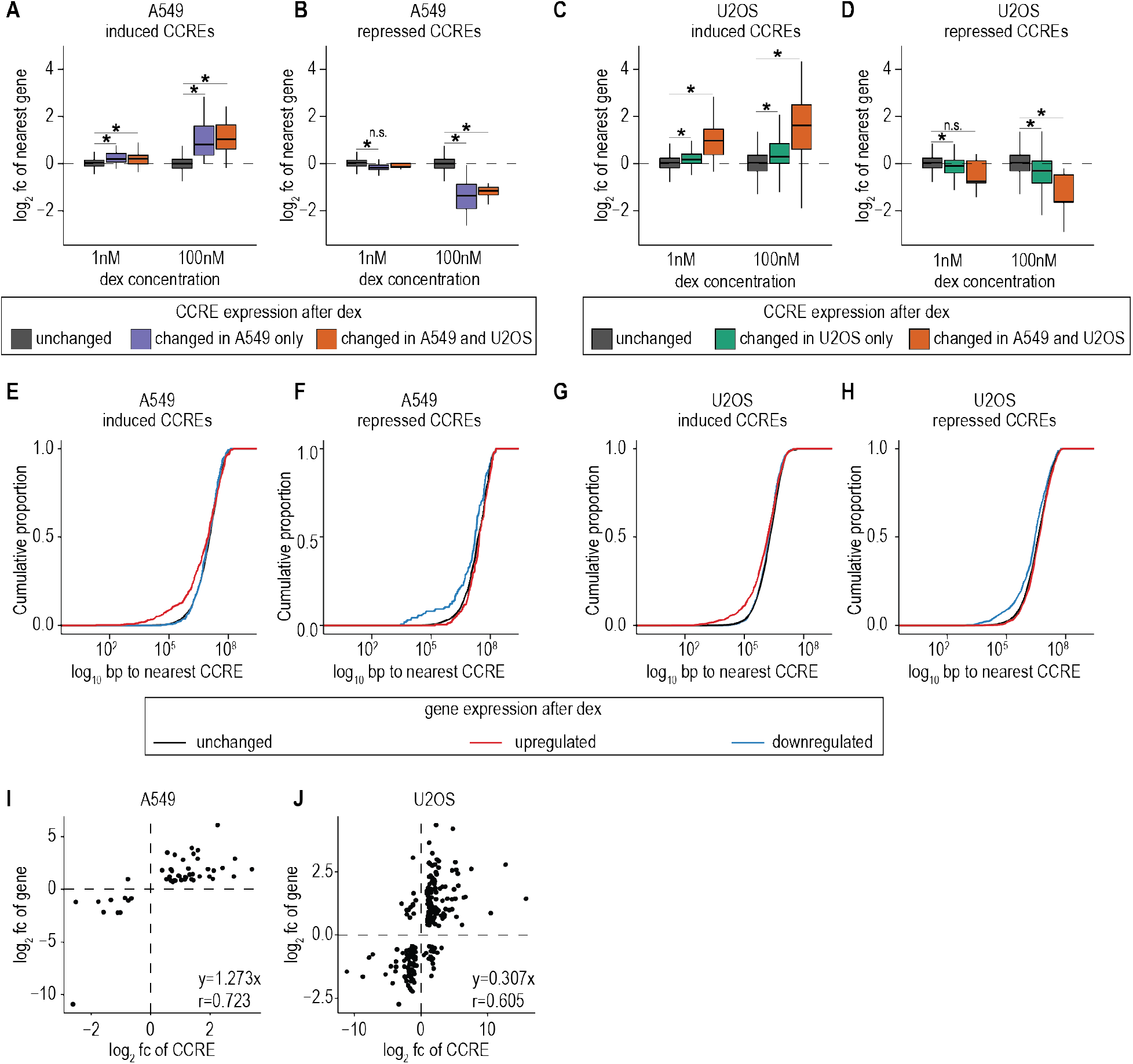
Enhancer characteristics present in CCREs identified by nascent transcription. **(A-D)** The closest annotated gene TSS for each differentially expressed CCRE was identified in A549 **(A, B)** and U2OS **(C, D)** cells. The expression changes for the associated genes are plotted for induced **(A, C)** and repressed **(B, D)** CCREs. Significance testing was performed with Wilcoxon rank sum tests; *p < 0.0005. **(E-H)** The closest induced **(E, G)** or repressed **(F, H)** CCRE to each gene was identified in A549 **(E, F)** and U2OS cells **(G, H)**, and the distances are plotted from the gene’s TSS. **(I)** The closest TSS of a differentially expressed gene was found for each differentially expressed CCRE in A549 cells. The expression changes for each gene and the nearest CCREs are plotted; n=48 genes. **(J)** The closest TSS of a differentially expressed gene was found for each differentially expressed CCRE in U2OS cells. The expression changes for each gene and the nearest CCREs are plotted; n=214 genes.

We next examined if degree of transcriptional changes at dex-responsive CCREs correlates with expression changes of the nearest dex-responsive gene. Sometimes multiple CCREs had the same nearest gene, and so for those, we used the sum of the CCREs’ induction signal. In both cell types, a significant correlation exists between the degree of transcriptional change for CCREs and for genes after dex treatment (Figure 4I,J). Interestingly, the slope of the best-fit line for gene and CCRE induction differ between the two cell types. In A549 cells (Figure 4I), the slope is close to one, showing that CCREs and their neighboring genes are similarly induced. In U2OS cells (Figure 4J), however, some CCREs have much stronger induction than their candidate target genes, implying that factors beyond transcription level affect enhancer activity on gene expression. Recent work shows that during macrophage differentiation, enhancers can act additively or synergistically (Choi et al. 2021), in alignment with our data. The overall correlation between CCRE transcription and nearby gene transcription is consistent with the idea that these CCREs enhance transcription of nearby target genes.

### GR binding is not sufficient for changes in CCRE transcription

Strikingly, <5% of GORs display significant changes in transcription in response to dex, whereas 67% of dex-induced CCREs are GORs (Figure 5A). Of the induced CCREs that lack GR binding, approximately 30% reside within GR-inducible genes, and thus their apparent induction could be an artifact of the gene’s increased transcription. Others may be induced due to long-range interactions, GR tethering, or indirect effects of the dex treatment. Only 3% of CCREs with repressed transcription are GR-bound (Figure 5A). PRO-seq profiles of CCREs generally show a bidirectional pattern of transcription, with TF motifs between the two transcription start sites (Core, Martins, et al. 2014; Tippens et al. 2020). We examined the PRO-seq signal at GORs with and without induced CCREs in A549 and U2OS cells (Figure 5B, C). These profiles are centered on the GR binding summit, and we observe the expected bidirectional pattern in induced CCREs, showing that although GR is not sufficient for transcription, GR binding between the two transcripts likely causes induction.

**Figure 5:**
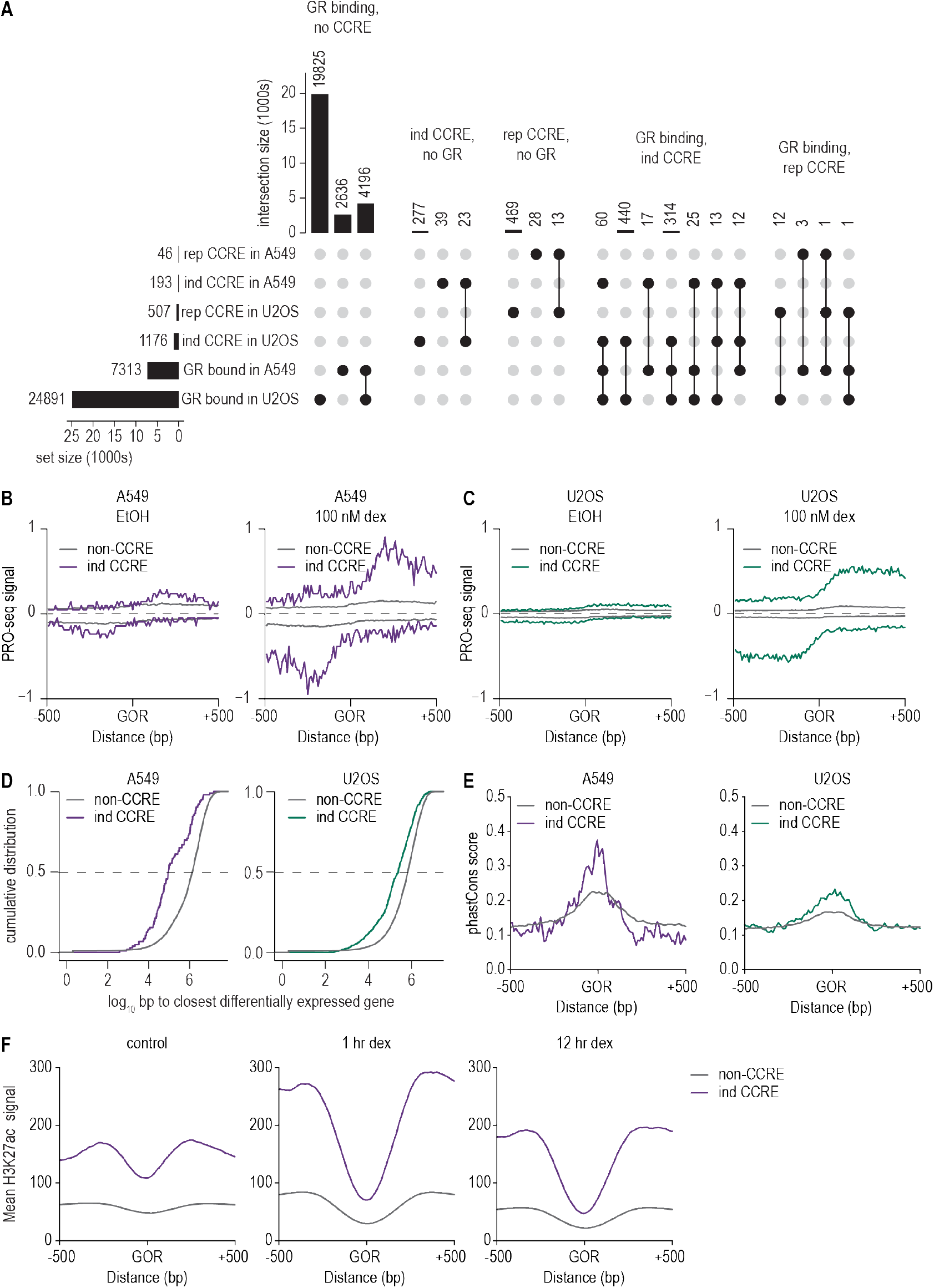
Most GR-bound sites do not have increased transcription. **(A)** UpSet plot showing overlap between GR binding and CCREs that are significantly induced (ind) or repressed (rep). **(B and C)** PRO-seq signal at GORs that have increased (“ind CCRE”) or no increase (“non-CCRE”) in transcription after 100 nM dex treatment in A549 **(B)** and U2OS **(C)** cells. **(D)** Cumulative distributions of distance between each GOR, based on the presence or absence of an induced CCRE, to the nearest induced gene. Significance testing by Wilcoxon rank sum tests; for both cell types p < 1 × 10^-15^. **(E)** Conservation at GORs with and without CCRE induction. **(F)** H3K27ac signal for GORs with and without CCRE induction.

We were interested in determining if GORs with and without changing CCRE transcription differed in other ways. GORs with increased transcription were significantly closer to induced genes than ones without changed transcription (Figure 5D) and had higher conservation scores within primates (Figure 5E), demonstrating that they are under purifying selection. These sites have higher H3K27ac signal in control cells that sharply increases after 1 hour of dex treatment (Figure 5F). H3K27ac is a chromatin modification thought to mark active enhancers (Creyghton et al. 2010). We considered that GORs without changing transcription at 45 minutes may gain enhancer activity later in the GR response. To address that hypothesis, we used existing H3K27ac data in A549 cells treated with dex for 12 hours (McDowell et al. 2018). We found that after 12 hours, GORs with no change in transcription after 45 minutes of dex have less H3K27ac than ones with induced transcription (Figure 5F). GORs with dex-responsive CCREs therefore have features that correlate with enhancer activity. While GORs without changes in transcription may have roles in the cellular response to GR, they appear to have less direct enhancer activity than ones with induced transcription.

We reasoned that the GORs with induced transcription would differ in TF composition from those without induced transcription. GORs with induced transcription in both A549 and U2OS cells were selectively enriched for FOSL2 motifs, and A549-specific GORs were also enriched for Ets motifs (Table S3). Binding of the AP-1 factor FOSL2 is also found at GORs that are present in both A549 and U2OS cells (Figure 2B), regardless of change in transcription. Of GORs bound in both cell types, 8% have increased transcription, as compared to 1% of A549-specific GORs and 2% of U2OS-specific GORs (Figure 5A). AP-1 motifs therefore may potentiate the GR response.

### Pioneer factor binding distinguishes A549- and U2OS-specific CCRE induction

We showed above that cell type-specific binding of GR corresponds to pioneer factor binding (Figure 2). Here, we determined if pioneer factor binding relates to the transcriptional response at GORs by comparing CEBPB and FOXA2 binding in non-dex treated A549 cells (Figure 6A, B). The loci at which GR binds specifically in U2OS cells show minimal binding of pioneer factors in A549 cells, whereas the intensity of CEBPB and FOXA2 binding is similar when comparing A549-specific GORs that do and do not overlap CCREs. These factors therefore appear to open chromatin and allow binding of other factors, but they are not sufficient for inducing transcription. As described above, AP-1 motifs mark the GORs that have induced transcription. In control A549 cells, the AP-1 component FOSL2 has greater binding to GORs that will have induced transcription upon dex addition (Figure 6C), suggesting that AP-1 poises these GORs for GR induction. Induced GR binding then follows the same pattern as JUNB binding (Figure 6D). We speculate from these findings that pioneer factors open chromatin, allowing AP-1 binding to mark the loci, which upon dex treatment enables GR binding and presumptive enhancer activity.

**Figure 6:**
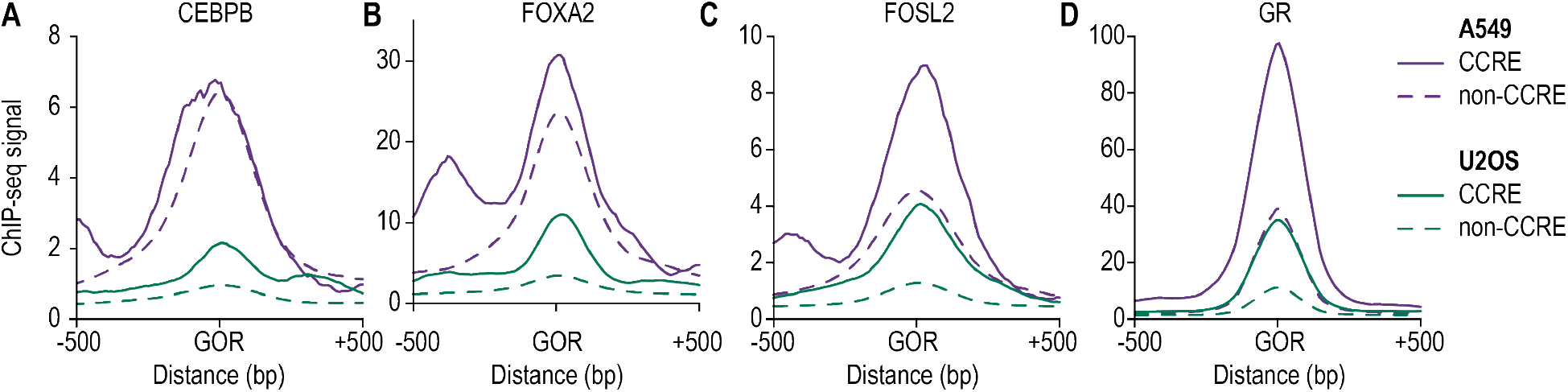
Pioneer factors underlie cell type-specific GR induction. ChIP-seq data for **(A)** CEBPB, **(B)** FOXA2, **(C)** FOSL2, and **(D)** GR from A549 cells are shown as metaplots centered on cell type-specific GORs of A549 and U2OS, with and without increased transcription.

## Discussion

We demonstrate here that cell type-specific GR responses require non-GR transcription factors, some of which are broadly expressed, and some which have limited expression. These classes of TFs assemble in a combinatorial manner, and none alone is sufficient for the cellular response to glucocorticoids. We primarily focused on the non-promoter genomic loci at which GR binds, as these make up >95% of GORs. Critically, we show that a small fraction of GORs has dex-responsive transcription in either cell type. Those responsive CCREs have binding sites and/or motifs for putative pioneer factors near the GR binding site, as well as for AP-1. Glucocorticoid-induced CCRE transcription therefore appears to require multiple classes of TFs, which have been implicated in other studies to affect distinct steps of the transcription cycle, including opening chromatin, maintaining accessibility, recruiting Pol II, and releasing Pol II into active elongation. Our study confirms and also expands upon previous work showing that a single stimulus acting on a single TF can regulate sets of genes that are the same in two cell types or specific to a particular cell type (Gertz et al. 2013; Weikum et al. 2017; Vihervaara, Duarte, et al. 2018)—likely a consequence of cell type-specific 3D chromatin architecture and of enhancers that are composed of different combinations of transcription factors. Importantly, it is likely that GR-mediated transcriptional regulation is conferred across the genome and in different cell types by a myriad of GR-containing transcriptional regulatory complexes, distinct in composition and conformation (Weikum et al. 2017), and displaying different combinations of characteristics that have been correlated with enhancer function.

Previous studies of GR signaling demonstrated roles for pioneer factor binding (So et al. 2007; Belikov et al. 2009; Grøntved et al. 2013; McDowell et al. 2018), pre-wired enhancer-promoter loops (D’Ippolito et al. 2018), and AP-1 binding (So et al. 2007; Biddie et al. 2011; Vockley et al. 2016; McDowell et al. 2018). Our critical addition was using PRO-seq to distinguish between GORs with and without dex-responsive nascent transcription, and to do so in two cell types that express different pioneer factors. GORs in A549 cells globally associate with the binding of FOXA2 and CEBPB pioneer factors (Figure 2). A small fraction of those GORs exhibit differential transcription after dex treatment that are distinguished by also displaying AP-1 binding (Figure 6). A subset of GORs is only present in U2OS cells. The corresponding loci in A549 cells lack FOXA2 and CEBPB binding and are less accessible, and so even though a GBS is present, the chromatin environment does not appear to be amenable to GR binding. Furthermore, AP-1 binding is seen at GR loci with dex-responsive CCRE transcription in A549 cells, and its motif is present at the transcribed GR loci in U2OS cells. AP-1 is therefore not essential for GR binding but may play a different role in regulating CCRE transcription. For example, AP-1 may maintain a larger region of accessible chromatin than FOXA or CEBP, thereby facilitating transcription at those sites. Alternatively, AP-1 may promote Pol II recruitment, poising the sites for transcription upon GR binding. GR has been reported to impact both Pol II recruitment and pausing at promoters (Gupte et al. 2013; Sacta et al. 2018), and putative enhancers can also undergo Pol II pausing (Core, Martins, et al. 2014; Scruggs et al. 2015; Henriques et al. 2018). Thus our comparative analysis of GR activity in two cell types identifies different roles for cell type-specific factors and for the more broadly expressed AP-1 TF.

While only a small fraction of GORs displays induced transcriptional activity after 45 minutes of dexamethasone treatment, others may be transcribed at later time points. This mechanism could occur because GR may itself decompact chromatin and recruit AP-1 binding (Vockley et al. 2016; Jubb et al. 2017). Direct validation of GR-mediated transcriptional regulation by CCREs (Ehmsen et al. 2019), together with high-resolution and extensive time course measuring nascent RNA, TF and coactivator binding, and chromatin accessibility, will further illuminate how GR and other factors coordinate to regulate gene expression.

Our work raises an interesting question: what is the role of GR’s widespread binding in the cellular response to glucocorticoids? GORs that exhibit changed transcription also include characteristics that correlate with enhancer activity. In particular, dex-responsive CCREs are close to dex-responsive genes. These sites also have higher conservation and increased H3K27ac after dex treatment. GR binds to thousands of other loci, however. Unequivocal determination of enhancer activity, and assessment of the role of transcription at those sites, would require extensive mutagenesis screening. We speculate that a subset of these GORs will undergo transcription later. Others could strengthen or modulate looping interactions in the genome and thus affect expression of genes distal from their enhancers. GR has also been implicated in condensate formation that increases the local concentration of transcription factors (Sabari et al. 2020; Garcia et al. 2021). The strong context specificity of GR action opens additional possibilities. GORs that are non-functional after dex treatment in A549 or U2OS cells could confer responses to a different glucocorticoid, in a different cell type, in combination with a particular kinases or other signal transduction machineries, at a specific developmental stage, or in a constrained three-dimensional tissue architecture. While failure to confer regulatory activity in one context is not an indication that a GOR is globally nonfunctional, it is possible that some GORs exist because TF motifs are degenerate and may form in the absence of selection.

The GR response has similarities to other inducible systems, including heat shock and TNF*α* treatment, showing that analysis of diverse systems of inducible gene expression can reveal common features of transcriptional regulation. In each of these instances, chromatin contacts between putative enhancers and promoters appear to be pre-wired (Jin et al. 2013; D’Ippolito et al. 2018; Ray et al. 2019). The fact that three different systems that all include a rapid transcriptional response to an extracellular stimulus already have the required 3D chromatin architecture begs the questions of how these contacts are specified during development and to what other stressors cells are prepared to respond. In Drosophila, many promoter-enhancer loops form early in development before the gene is activated, and the promoters typically display polymerase pausing (Ghavi-Helm et al. 2014), which likely sets these genes up to be rapidly induced when needed. Further, GR, HSF1, and NF-*κ*B (the TF that directs the TNF*α* response) bind to many sites that fail to rapidly increase transcription in certain contexts (Jin et al. 2013; Mahat, Salamanca, et al. 2016; Vockley et al. 2016; Vihervaara, Mahat, et al. 2017). This TF binding without activity suggests that different contexts, including the presence and activity of other factors, are required for transcription. Both HSF1 and NF-*κ*B act by releasing paused Pol II (Barboric et al. 2001; Danko, Hah, et al. 2013; Duarte et al. 2016), so the presence of other factors that facilitate Pol II recruitment and initiation may be essential (Duarte et al. 2016). For GR, AP-1 binding may commonly play a role in GR-mediated induction of transcription, and more work is needed to define mechanisms by which AP-1 acts. These studies broadly show that transcription is regulated at multiple steps, all of which need to act in concert for the appropriate cellular response to hormone signaling.

## Methods

### Cell culture

U2OS and A549 human-derived parental cell lines were obtained from American Type Culture Collection and authenticated by short tandem repeat (STR) profiling (ATCC); U2OS.hGR cells were generated previously in the Yamamoto laboratory (Rogatsky, Trowbridge, et al. 1997). Cells were maintained in DMEM/low glucose (4 mM L-glutamine, 1000 mg/L glucose, 110 mg/L sodium pyruvate; HyClone Laboratories, Inc.) supplemented with 5% FBS (Gemini BioProducts) at 37°C with 5% CO2.

### PRO-seq library generation

A549 and U2OS.hGR cells were treated with ethanol (Koptek) or 100 nM dexamethasone (Sigma) for 45 minutes. PRO-seq libraries were generated as described in (Mahat, Kwak, et al. 2016). Approximately 8 × 10^6^ cells were collected by scraping in 1x PBS supplemented with 10 mM EDTA on ice. All spins were performed in a swinging bucket rotor at 4°C at 1000 g for 5 minutes. After pelleting, cells were then washed twice with 1x PBS. They were incubated in 5 mL permeabilization buffer (10 mM Tris-HCl, pH 7.4; 300 mM sucrose; 10 mM KCl; 5 mM MgCl2; 1 mM EGTA; 0.05% Tween-20; 0.1% NP40 substitute; 0.5 mM DTT, 1x Pierce Protease Inhibitor Tablet (ThermoFisher); 4 units per mL SUPERaseIn RNase Inhibitor (ThermoFisher)) for 5 minutes on ice, then washed twice in permeabilization buffer. Cells were resuspended in 100 *μ*L storage buffer (50 mM Tris, pH 8.0; 25% glycerol (v/v), 5 mM MgAc2, 0.1 mM EDTA, 5 mM DTT, 40 units per ml SUPERaseIn RNase Inhibitor (ThermoFisher)), then flash frozen in liquid nitrogen and stored at −80°C.

The run-on reaction mixture was made with biotinylated rCTP and rUTP (5 mM Tris, pH=8.0; 2.5 mM MgCl2; 0.5 mM DTT; 150 mM KCl; 200 *μ*M rATP; 200 *μ*M rGTP; 20 *μ*M Biotin-11-rCTP (PerkinElmer); 20 *μ*M Biotin-11-rUTP (PerkinElmer); 0.8 units per *μ*L SUPERaseIn RNase Inhibitor (ThermoFisher); 50% sarkosyl), then preheated at 37°C for at least 5 minutes. 100 *μ*L of run-on mixture was added to 100 *μ*L of thawed permeabilized cells, then incubated at 37°C for 5 minutes. The reaction was stopped by adding Trizol LS (ThermoFisher), followed by RNA isolation. RNA was fragmented on ice for 8 minutes with 0.2 N NaOH and neutralized with Tris pH 6.8, followed by buffer exchange with a Bio-Gel P-30 column (Bio-Rad).

Nascent RNA was enriched with M-280 Streptavidin Dynabeads (ThermoFisher), then Trizol extracted. In-line barcoded 3′ adapters were ligated with T4 RNA ligase I (NEB) for 12 hours at 20°C, followed by a second enrichment with M-280 Streptavidin Dynabeads (ThermoFisher) and Trizol extraction. The 5′ end was repaired by decapping with RppH (NEB) for 1 hour at 37°C, then phosphorylated with T4 PNK (NEB) for 1 hour at 37°C. The 5′ adapter, which included a 6-nucleotide UMI, was ligated with T4 RNA ligase I (NEB) for 12 hours at 20°C, followed by a third enrichment with M-280 Streptavidin Dynabeads (ThermoFisher) and Trizol extraction. cDNA was generated with SuperScript III (ThermoFisher), and libraries were amplified with Phusion for 13 cycles. Libraries were cleaned up with Ampure XP beads (Beckman Coulter) at a 1:1.6 ratio.

### PRO-seq sequencing

Libraries were sequenced on a HiSeq 4000 with single-end 50 nt reads. Libraries were separated based on their in-line barcodes with barcode_splitter v0.18.0 (Leach, Parsons 2019), PCR duplicates were removed with prinseq v0.20.4 (Schmieder, Edwards 2011), and adapters were trimmed with cutadapt v1.16 (Martin 2011). Reads were aligned to hg38 with bowtie2 v2.3.4.1 (Langmead, Salzberg 2012), using end-to-end alignment and the sensitive settings. Uniquely mapping reads were subsampled to the sample with the lowest read number with samtools v1.7. Bedgraphs were generated in which the read numbers at the 3′ ends were reported using bedtools v2.26.0 (Quinlan, Hall 2010), then converted to bigwigs (WJ Kent et al. 2010). Data were visualized in the Integrated Genome Viewer (Robinson et al. 2011).

### PRO-seq data analysis

Transcriptional regulatory elements (TREs) for each cell type and treatment were identified using dREG Gateway (Danko, Hyland, et al. 2015; Z Wang et al. 2019; Chu et al. 2019) on subsampled data sets. Genes were considered potentially expressed if a TRE overlapped at least one annotated RefSeq transcription start site (TSS). TREs that were at least 1 kb away from an annotated (TSS) were considered candidate *cis*-regulatory elements (CCREs), and a unified set of CCREs were determined by merging CCREs from each cell type and treatment, using bedtools v2.26.0 (Quinlan, Hall 2010).

Read counts for genes and CCREs were found using the R v3.6 (R Core Team 2019) package bigwig (Martins, Kent 2014). Differential expression and log2 fold-change values were determined with DESeq2 (Love, Huber, et al. 2014), using a Benjamini-Hochberg-adjusted p-value cut-off of 0.05 as significantly differentially expressed. k-means clustering was performed with the PAM package in R, and differential expression data were plotted with the R packages ggplot2 (Wickham 2016) and pheatmap.

Motif enrichment was performed with HOMER v3.12 (Heinz et al. 2010) to find differentially enriched motifs, using 300 nt segments centered on the TRE, with all CCREs as the background set. The most enriched motifs were compared to HOCOMOCO v11 (Kulakovskiy et al. 2018).

For analyses concerning expression of genes near CCREs, the gene set used was filtered to expressed genes, which were defined as having an annotated TSS overlapping a site of bidirectional transcription, as called from dREG. In Figure 3B, all expressed genes were used, whereas in Figure 3C, only ones that were significantly differentially expressed (Benjamini-Hochberg p-value < 0.05, log2 fold-change > 0) were included. The CCREs used in Figure 3C were required to be intergenic and within 100 kb of the closest dex-responsive gene. The TSS closest to each CCRE was found using bedtools closest (Quinlan, Hall 2010). Scripts used for these analyses are available at https://github.com/ewissink/GR-pioneer-factors-AP1-enhancers.

### GR ChIP-seq

A549 and U2OS.hGR cells were treated for 90 minutes with dexamethasone (Sigma) at 100 nM or ethanol (Koptec), harvested by trypsinization, and fixed in suspension by adding 36.5% formaldehyde to a final concentration of 1% v/v; after incubating 3-10 minutes at room temperature (RT), formaldehyde was quenched by adding 2.5 M glycine to 0.3 M, RT 5 minutes, followed by transfer to ice. After washing twice in 20 mL ice-cold TBS (100 mM Tris-HCl, pH 7.5 @ 4°C/150 mM NaCl, centrifugation at 450g, 5 minutes, 4°C), cells were washed 3x in 1 mL RT MC lysis buffer (10 mM Tris-HCl, pH 7.5 @ RT, 10 mM NaCl, 3 mM MgCl2, 0.5% (v/v) Tergitol type NP-40), resuspended in 1 mL RT MC lysis buffer, pelleted at 200g, 5 minutes, and frozen in liquid N2 for storage at −80°C.

Frozen nuclear pellets were resuspended in 180 *μ*L MNase reaction buffer (10 mM Tris-HCl, pH 7.5 @ RT, 10 mM NaCl, 3 mM MgCl2, 1 mM CaCl2, 4% (v/v) Tergitol type NP-40) supplemented with PMSF to 1:100, and the volume was adjusted to 270 *μ*L with MNase reaction buffer. 1.35 *μ*L MNase (New England Biolabs, diluted 1:10 in MNase reaction buffer) was added to the resuspended chromatin and incubated at 37°C, 5 minutes. A cOmplete Mini, EDTA-free Protease Inhibitor Cocktail tablet (Roche) was dissolved in 500 *μ*L MNase buffer (PIn cocktail); MNase reaction was stopped by adding 5.4 *μ*L 0.2 M EGTA (pH 8), 7.2 *μ*L 100 mM PMSF, 14.65 *μ*L PIn cocktail, 14.65 *μ*L 20% SDS, and 14.4 *μ*L 5 M NaCl, with gentle tube inversion to mix. 164 *μ*L volume was transferred to each of two 1.5 mL Bioruptor^®^ Plus TPX microtubes (Diagenode, Denville, NJ) and sonicated in a Bioruptor^®^ Plus (UCD-300) Sonication System using intensity setting ‘H’ (320 W) and sonication parameters “CYCLE Num:30, Time ON:30sec, Time OFF:30sec”. During a 15-minute rest interval, samples were vortexed for 5 seconds, spun down in a microfuge, and transferred to new TPX tubes on ice, followed by a second round of sonication (amounting to 60 cycles total).

For GR ChIP, 150 *μ*L Dynabeads™ Protein G slurry (Invitrogen) was mixed with 20 mg N499 (rabbit *α*-human GR IgG) antibody (Rogatsky, Zarember, et al. 2001) plus 450 *μ*L Lysis Buffer 2 (10 mM Tris-HCl, 1 mM EDTA, 150 mM NaCl, 5% (v/v) glycerol, 0.1% (w/v) sodium deoxycholate, 0.1% (w/v) SDS, 1% (v/v) Triton X-100, pH 8 @ 4°C; no PIn added) in a 1.5 mL Eppendorf tube, incubated 1 hour with rolling in 4°C cold room; tubes were placed in magnetic rack and supernatant was removed. Sonicated chromatin samples were pelleted at maximum speed, 10 minutes, 4°C, and white pellet and cloudy suspension above pellet were recovered by transferring from 1.5 *μ*L TPX tubes to new 1.5 mL tubes. 10 *μ*L aliquot of combined input was set aside for later processing. 25 *μ*L 100X Halt™ Protease Inhibitor Cocktail (Thermo Scientific) was added to a 5 mL tube with 2.5 mL Dilution Buffer (identical to Lysis Buffer 2, except without SDS). 275 *μ*L chromatin (from approximately 1.5 × 10^7^ cells) was added to the Dilution Buffer+beads in the 5 mL tube, effectively diluting the chromatin 1:10. The 5 mL tube was sealed with parafilm and incubated 4 hours on a roller in a 4°C cold room. During this time, input material was reverse-crosslinked by adding TE (10 mM Tris-HCl, 1 mM EDTA, pH 8; pH 7.5 @ RT) to a total volume of 80 μL, followed by addition of 100 *μ*L ChIP Elution Buffer (50 mM Tris-HCl, pH 7.5 @ RT, 10 mM EDTA, 1% SDS) and 20 *μ*L Pronase (Roche, 20 mg/mL) for incubation at 42°C for 2 hours, then 65°C overnight.

100X Halt PIn was warmed to RT, and 15 *μ*L was added to 1.5 mL of each of three Wash Buffers (A-C; Wash Buffer A: 10 mM Tris-HCl, 1 mM EDTA, 150 mM NaCl, 5% (v/v) glycerol, 0.1% (w/v) sodium deoxycholate, 0.1% (w/v) SDS, 1% (v/v) Triton X-100, pH 8.0 with Halt protease inhibitor mixture; Wash Buffer B: Buffer A with 500 mM NaCl and Halt protease inhibitor mixture; Wash Buffer C: 20 mM Tris-HCl, 1 mM EDTA, 250 mM LiCl, 0.5% (v/v) Nonidet P-40, 0.5% (w/v) sodium deoxycholate, Halt protease inhibitor mixture, pH 8.0). Beads were washed consecutively in 1.5 mL of each of the Wash Buffers A-C (1X Halt PIn), by gently adding buffer to resuspend the beads, then placing the tube on a magnetic rack and removing the supernatant after beads settled. Chromatin (from 1.5 × 10^7^ cells on beads from 150 *μ*L slurry) was eluted from beads in 300 *μ*L Elution Buffer/Reverse-Crosslinking Buffer (10 mM Tris-HCl, 1 mM EDTA, 0.7% (w/v) SDS, pH 8 @ RT) by incubating beads in buffer for 5 minutes, RT with gentle pipetting to occasionally mix, then allowing beads to settle on magnetic rack and transferring eluant volume to a new 1.5 mL Eppendorf tube. To reverse crosslinks, 450 *μ*L Adjustment Buffer (50 mM Tris, 10 mM EDTA, 0.45% SDS pH 7.0 at RT) was added with 82.5 *μ*L Pronase (20 mg/mL) to 300 *μ*L eluted chromatin, followed by incubation at 42°C for 2 hours, then 65°C overnight. DNA was subsequently cleaned from “input/MNase only” samples using a Qiagen PCR Purification Kit, and MinElute PCR Purification Kit (Qiagen) columns were used to purify ChIPs (one column/each ChIP, using 2.5 mL ERC buffer or 4.16 mL PB buffer), eluted in 15 *μ*L EB. Recovered DNA was stored at −20°C. Libraries were generated using an Ovation^®^ Ultralow System V2-32 (NuGEN Technologies, Redwood City, CA), quantified on a 2100 Bioanalyzer System (Agilent, Santa Clara, CA) with High Sensitivity DNA Kit.

### ChIP-seq sequencing

GR ChIP-seq libraries were sequenced on a HiSeq 4000 with single-end 50 nt reads. The analysis was performed with Galaxy (Afgan et al. 2018). Reads were aligned to hg38 with bowtie v1.1.2 (Langmead, Trapnell, et al. 2009), using the setting –best and with one alignment reported per read. PCR duplicates were removed with samtools v1.3.1 rmdup. Peaks were called with MAC2 v2.1.1 (Zhang et al. 2008), using the input as the control file. Bedgraphs of the q values were generated with MACS2 and used for downstream analyses.

### ChIP- and ATAC-seq analyses

Fastq files for ChIP-seq and ATAC-seq datasets that were generated by other lab groups were downloaded from ENCODE or SRA, as described below. Reads were aligned to hg38 with bowtie2 (Langmead, Salzberg 2012). Signal tracks were produced using MACS2 (Zhang et al. 2008), using ENCODE pipelines (Moore et al. 2020) (https://github.com/ENCODE-DCC/atac-seq-pipeline; https://github.com/ENCODE-DCC/chip-seq-pipeline). Data were visualized in the Integrated Genome Viewer (Robinson et al. 2011). Heatmaps and metaplots were generated with Deeptools v3.4.3 (Ramírez et al. 2016).

Additional datasets used. The following data sets were used in this study: U2OS.GR ATAC-seq, SRA: (BH Lee, Stallcup 2018); A549 ATAC-seq, ENCODE: ENCSR220ASC (https://www.encodeproject.org/) (ENCODE Project Consortium 2012; Davis et al. 2018); A549 FOXA2 ChIP-seq, SRA: GSE90454 (Donaghey et al. 2018); A549 CEBPB ChIP-seq, ENCODE: ENCSR701TCU (McDowell et al. 2018); A549 H3K27ac ChIP-seq, ENCODE: ENCSR778NQS (McDowell et al. 2018); A549 JUNB ChIP-seq, ENCODE: ENCSR656VWZ (McDowell et al. 2018), A549 FOSL2 ChIP-seq, ENCODE: ENCSR593DGU (McDowell et al. 2018); A549 JUN ChIP-seq, ENCODE: ENCSR192PBJ (McDowell et al. 2018). PhastCons scores for 30 primates were accessed from the UCSC Santa Cruz Genome browser (Pollard et al. 2010).

## DATA ACCESS

All raw and processed sequencing data generated in this study have been submitted to the NCBI Gene Expression Omnibus (GEO; https://www.ncbi.nlm.nih.gov/geo/) under accession numbers https://www.ncbi.nlm.nih.gov/geo/query/acc.cgi?acc=GSE168767 (PRO-seq) and GSE163398 (ChIP-seq).

## Acknowledgments

This work was supported by the NCI grant R01-CA020535 (to K.R.Y.), the NHGRI grant UM1-HG009393 (to J.T.L.), and the NIGMS grant F32-GM129904 (to E.M.W.). We thank the UCSF Center for Advanced Technology for high-throughput sequencing assistance. We also thank Charles Danko, the Lis lab, and Yamamoto lab for invaluable discussions. This LaTeX template was provided by George Kour.

