## Supplementary Figures for "Glucocorticoid receptor collaborates with pioneer factors and AP-1 to execute genome-wide regulation"

**Supplementary Material**

Table S1: Gene expression data from DESeq2

Table S2: MACS2 output for GR ChIP-seq

Table S3: HOMER output for motif searching

Table S4: CCRE expression data from DESeq2

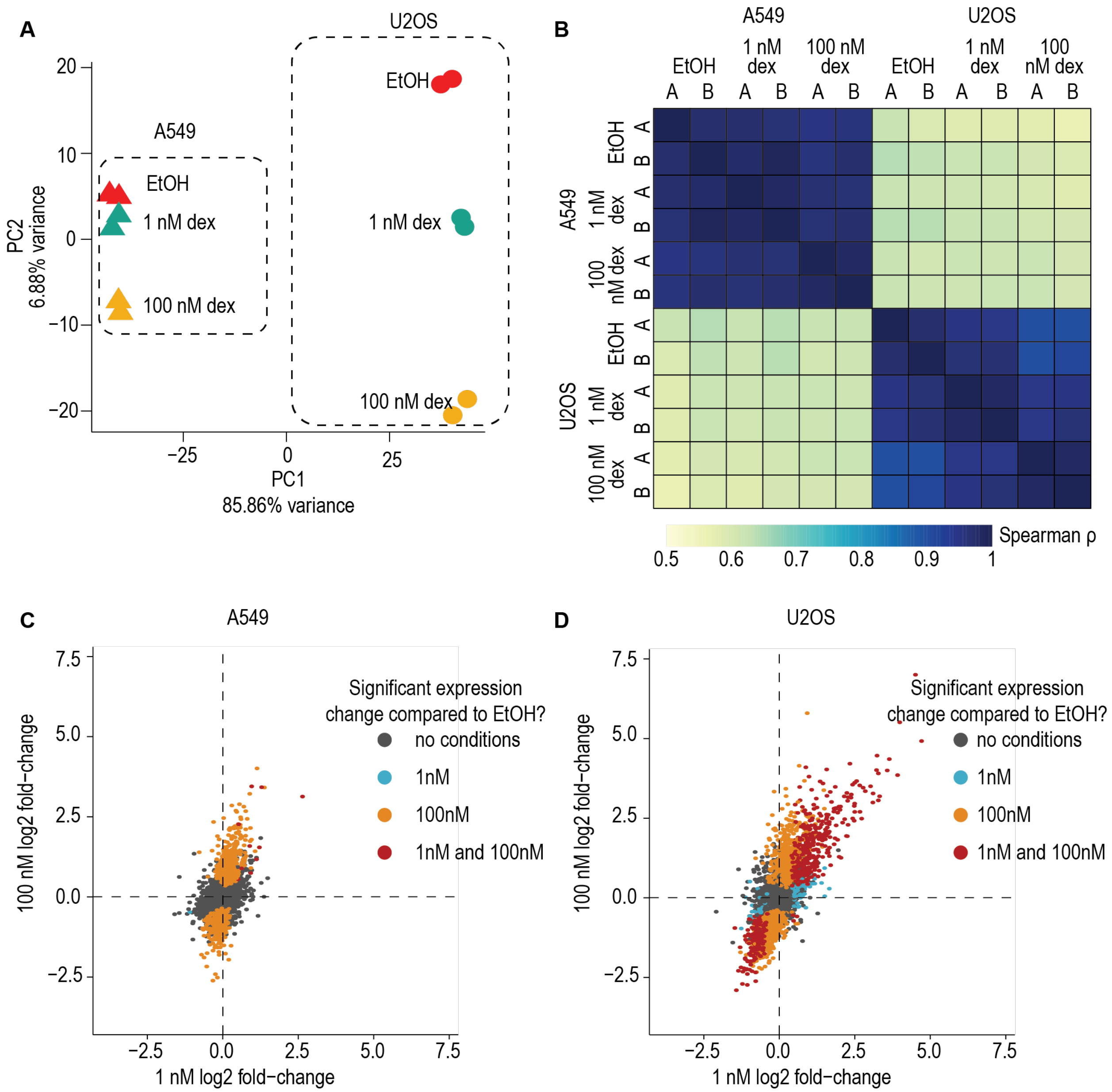


**Figure S1, related to Figure 1. (A)** PCA of gene expression in A549 and U2OS cells after dex treatment. Cell types separate on the x-axis, dex dosages separate on the y-axis. **(B)** Spearman correlation coefficients for gene expression in A549 and U2OS cells after dex treatment from PRO-seq data. Two biological replicates (A and B) were performed for each treatment (EtOH, 1 nM dex, and 100 nM dex) in the two cell types. Replicates correlated well, whereas A549 and U2OS cells have very different expression profiles from each other. **(C)** Expression change of genes in A549 cells after 1 nM and 100 nM dex treatment. Genes with significant changes in expression have Benjamini-adjusted $p < 0.05$. Correlation between all genes is Spearman $\rho$ = 0.499, $p<10^{-15}$. **(D)** Expression change of genes in U2OS cells after 1 nM and 100 nM dex treatment. Genes with significant changes in expression have Benjamini-adjusted $p < 0.05$. Correlation between all genes is Spearman $\rho$ = 0.705, $p<10^{-15}$.

**A**

| | log2(U2OS/A549) | FDR |
|---|---|---|
| FOXA1 | -4.6484210 | 1.054854e-37 |
| FOXA2 | -3.0651540 | 1.670134e-23 |
| CEBPA | -2.9754384 | 6.339162e-28 |
| CEBPB | -1.1932576 | 2.737302e-06 |
| RUNX1 | 0.8562571 | 1.198145e-07 |
| RUNX2 | 2.4861709 | 3.896162e-32 |
| NR3C1 (GR) | -1.1105145 | 4.151186e-10 |
| FOS | 0.7493683 | 5.931906e-02 |
| FOSL1 | 0.9151102 | 8.624001e-04 |
| FOSL2 | -0.9578456 | 1.658272e-09 |
| JUN | 0.5587163 | 3.311179e-01 |
| JUNB | -0.6966183 | 9.194504e-03 |

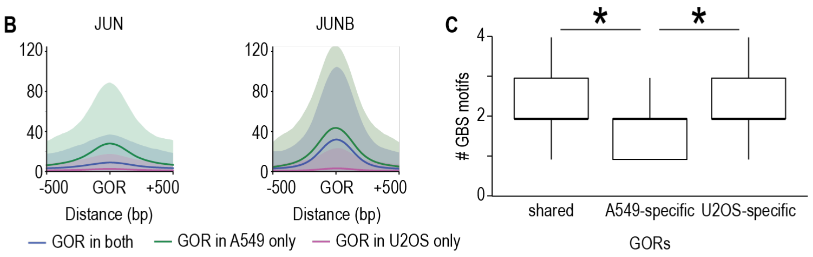


**D**

| cell type | change in gene exp | observed genes | expected genes | enrichment | Fisher test p value |
|---|---|---|---|---|---|
| A549 | induced | 76 | 16.1 | 4.72 | 1.452e-32 |
| A549 | repressed | 13 | 9.7 | 1.33 | 0.062 |
| U2OS | induced | 234 | 86.2 | 2.71 | 2.72e-60 |
| U2OS | repressed | 61 | 91.4 | 0.67 | 1.0 |

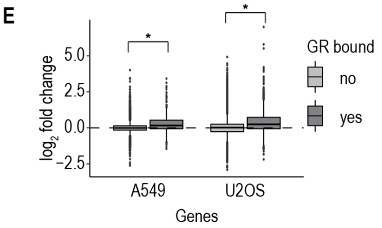


**Figure S2, related to Figure 2. (A)** Gene expression differences from DESeq2 for transcription factors that have motif enrichment in A549 or U2OS cells (Table S1). FOXA1, FOXA2, CEBPA, and CEBPB are more highly expressed in A549 cells. RUNX1 and RUNX2 are more highly expressed in U2OS cells. NR3C1, which is the gene symbol for GR, is more highly expressed in A549 cells. FOS, FOSL1, FOSL2, JUN, and JUNB are all components of AP-1, and some are more highly expressed in A549 cells, while others are more highly expressed in U2OS cells. **(B)** In A549 cells, metaplots of mean signal for JUN and JUNB in control cells are shown for shared GORs (blue), A549-specific GORs (green), and U2OS-specific GORs (pink). Signal is the -log10 p-value of enrichment, and shading indicates the standard deviation. **(C)** Number of GBS motifs present in GORs. *$p=2.2*10^{-16}$. **(D)** Enrichment of GR binding to promoters of induced and repressed genes in A549 and U2OS cells. Expected genes were calculated as (# genes with GR promoter binding/total # genes) * (# differentially expressed genes). **(E)** Comparison of log2-fold change in expression for genes without and with GR binding at their promoters. * $p < 10^{-8}$.

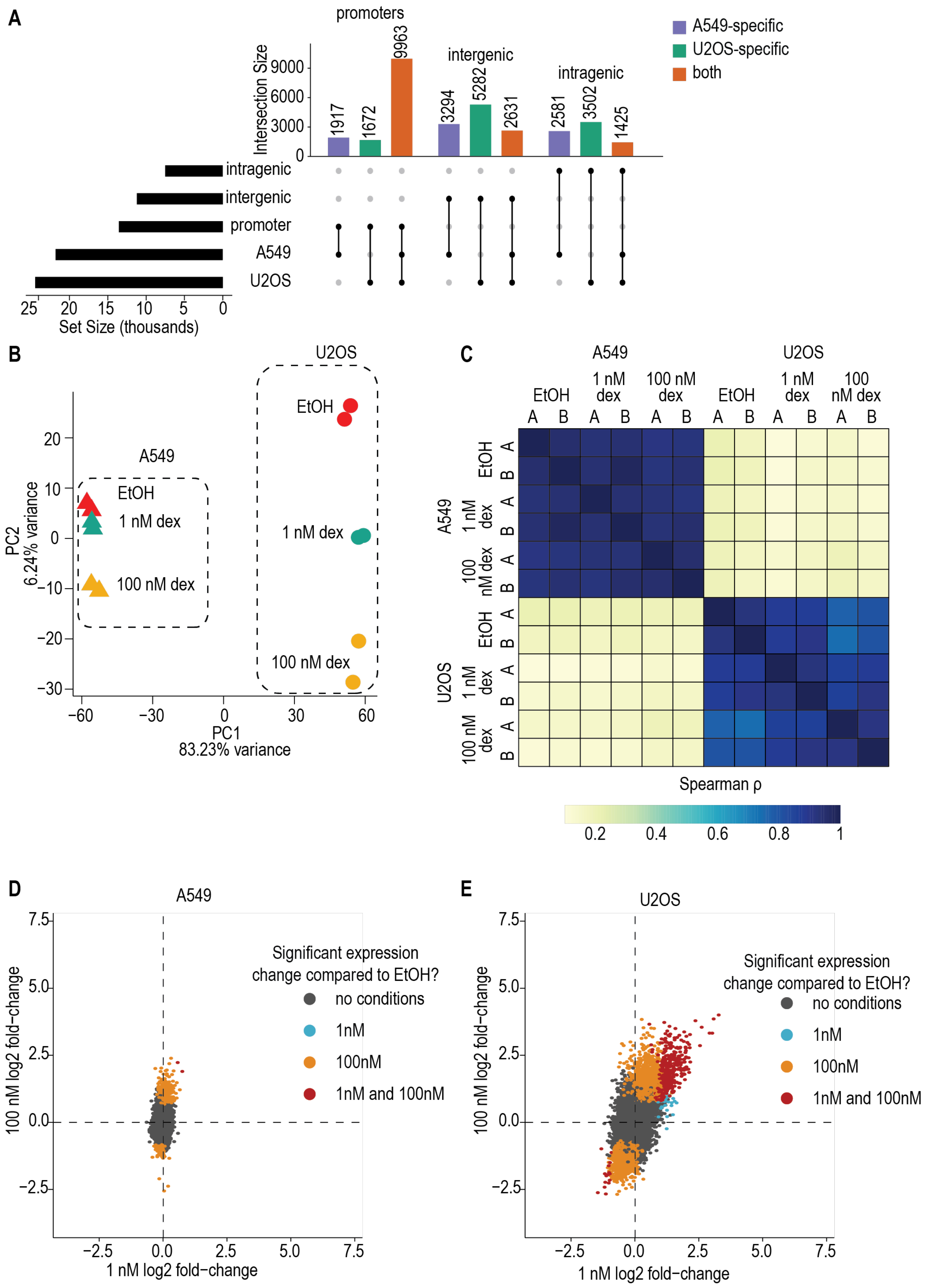
A
promoters
intergenic
intragenic
Intersection Size
A549-specific
U2OS-specific
both
Set Size (thousands)
B
U2OS
A549
EtOH
1 nM dex
100 nM dex
PC1
83.23% variance
PC2
6.24% variance
C
Spearman ρ
D
A549
Significant expression change compared to EtOH?
no conditions
1nM
100nM
1nM and 100nM
100 nM log2 fold−change
1 nM log2 fold−change
E
U2OS

**Figure S3, related to Figure 3.** **(A)** UpSet plot showing genomic distribution of dREG-called bidirectional transcripts, which consist of promoters and CCREs (which can be intergenic or intragenic). Their cell type-specificity is also shown. **(B)** PCA of CCRE expression in A549 and U2OS cells after dex treatment. Cell types separate on the x-axis, dex dosages separate on the y-axis. **(C)** Spearman correlation coefficients for CCRE expression in A549 and U2OS cells after dex treatment from PRO-seq data. Two biological replicates (A and B) were performed for each treatment (EtOH, 1 nM dex, and 100 nM dex) in the two cell types. Replicates correlated well, whereas A549 and U2OS cells have very different CCRE expression profiles from each other. **(D)** CCREs with significant changes in expression have Benjamini-adjusted $p < 0.05$. Correlation between all CCREs is Spearman $\rho = -0.055$, $p<10^{-11}$. **(E)** Expression change of CCREs in U2OS cells after 1 nM and 100 nM dex treatment. CCREs with significant changes in expression have Benjamini-adjusted $p < 0.05$. Correlation between all genes is Spearman $\rho = 0.440$, $p<10^{-15}$.
